# Reactivation of DRP1 plays a functional role in resistance to MEK inhibition in pancreatic cancer cells

**DOI:** 10.64898/2026.05.20.726663

**Authors:** Salma Sharmin, Jennifer A. Kashatus, Sara J. Adair, Emma Bakall Loewgren, Mohammad Fallahi-Sichani, Todd W. Bauer, David F. Kashatus

## Abstract

**Background:** In RAS-mutant tumors, ERK phosphorylates the mitochondrial fission GTPase DRP1 to promote mitochondrial fission. DRP1 activity is tumor-promoting in pancreatic and other RAS-driven cancers, but its role in therapeutic resistance is unknown.

**Methods:** We developed a panel of patient-derived pancreatic cancer cell lines resistant to the MEK inhibitor trametinib. We used immunofluorescence imaging, in vitro growth assays and orthotopic xenografts to determine the role of DRP1 in trametinib resistance.

**Results:** We find that trametinib-resistant cells exhibit increased expression and phosphorylation of DRP1 compared to sensitive counterparts. Quantitative analysis of mitochondrial structure reveals that mitochondria in resistant cells are morphologically distinct and relatively smaller than sensitive cells treated with trametinib. Genetic and pharmacological inhibition of both c-Myc and CDK6 are sufficient to block DRP1 phosphorylation in resistant cells, suggesting that activation of a c-Myc-CDK6 signaling axis drives reactivation of mitochondrial fission in the absence of MAPK signaling. Importantly, deletion of DRP1 leads to either growth inhibition or re-sensitization to trametinib in resistant lines.

**Conclusion:** These findings suggest DRP1 contributes to drug resistance, and that inhibition of mitochondrial fission might be a promising therapeutic strategy to combat resistance to MAPK and RAS inhibitors.

## Background

Pancreatic ductal adenocarcinoma (PDAC) patients have a 5-year survival rate of 13%, ranking it the third most lethal cancer in the United States [1,2]. Such dismal survival is due to the scarcity of effective treatments at the time of diagnosis and emphasizes the need to develop novel therapeutics. Greater than 90% of PDAC tumors harbor an activating mutation in KRAS which leads to engagement of downstream effector pathways that drive the tumorigenic phenotype [3]. Despite the development of potent inhibitors that target the key KRAS effector pathways, including the RAF-MEK-ERK MAPK pathway [4], and more recent development of inhibitors that target KRAS directly [5–7], targeted therapies have been largely unsuccessful in PDAC. The potential therapeutic value in targeting the RAS-MAPK axis in pancreatic cancer is evident from other tumor types, including melanoma, where direct targeting of mutant BRAF either alone, or in combination with MEK, improves overall survival [8–10] as well as non-small cell lung cancer, where direct targeting of mutant KRAS leads to improved overall survival [11]. However, direct inhibition of MEK with the drug trametinib did not improve survival of pancreatic cancer patients despite the well-established importance of MAPK signaling to support PDAC tumor growth [12]. These outcomes suggest a rapid development of resistance against RAS-MAPK inhibition in PDAC. Further understanding of resistance mechanisms developed within these tumors will be important to allow for the development of effective new treatment strategies.

Mitochondria in eukaryotic cells exhibit dynamic morphologies ranging from highly fragmented to highly interconnected [13]. The mitochondrial morphology of a cell is dictated by the relative activities of the dynamin-related GTPases DRP1, which regulates mitochondrial fission and MFN1/2 and OPA1, which regulate mitochondrial outer and inner membrane fusion, respectively. We and others have demonstrated that signaling downstream of oncogenic RAS results in a fragmented mitochondrial phenotype through ERK-mediated phosphorylation of DRP1 on serine 616 and a subsequent shift in the balance of fusion and fission activity [14,15]. Blocking DRP1 phosphorylation through inhibition of RAF or MEK shifts mitochondria to a more interconnected phenotype. Importantly, genetic inhibition of DRP1 decreases tumor burden and increases survival in two complementary in vivo models of KRAS-driven pancreatic cancer [14,16]. This work is consistent with the mitochondrial morphologies observed in other tumor types, including lung, breast, colon and brain [14,17–20] and the pro-tumorigenic functions that have been associated with cells exhibiting high relative mitochondria fission, including cell proliferation, stemness, immune escape, migration and invasion [18,21–23].

The growing importance of mitochondrial fission in tumorigenesis has led several groups to explore its relationship with chemoresistance. For example, mitochondrial fission contributes to cisplatin resistance in hypoxic ovarian cancer cells [24]. Additionally, high-mobility group box 1 (HMGB1) expression leads to oxaliplatin resistance in colorectal cancer cells through ERK-mediated DRP1 phosphorylation [25]. However, the role of DRP1 and mitochondrial fission in the development of resistance to therapies targeting RAS-MAPK signaling is unknown. To address this, we set out to dissect whether DRP1-mediated mitochondrial fission plays a role in acquired resistance to the MEK inhibitor trametinib in a panel of patient-derived pancreatic cancer cell lines. Interestingly, we find that DRP1 S616 phosphorylation is increased in pancreatic cancer cells resistant to trametinib and these cells exhibit distinct mitochondrial morphologies compared to sensitive cells. Further, we find that DRP1 S616 phosphorylation in trametinib-resistant cells is independent of ERK activity but is blocked following inhibition of the cyclin-dependent kinases CDK4 and CDK6 and the transcription factor c-Myc. Importantly, inhibition of either DRP1 or CDK4/6 inhibits proliferation of trametinib-resistant PDAC cell lines, indicating that this pathway may represent an important therapeutic target to prevent or overcome resistance to therapies targeting RAS-MAPK signaling.

## Methods

### Cell culture

Human pancreatic patient-derived xenograft cell lines 188 and 395 [16,26] and established patient-derived pancreatic cancer cell lines PANC-1 and CFPAC-1 (ATCC, VA, USA) were cultured in RPMI 1640 (Gibco #11875093, USA) supplemented with 10% FBS (Gibco #16000-044) and 1% penicillin/streptomycin (Gibco #15140-122). Cells were cultured at 37⁰C in a humidified incubator with 5% CO2. HEK-293T cells were cultured in DMEM (Gibco #11965-092) supplemented with 10% FBS (Gibco) and 1% penicillin/streptomycin (Gibco). Drugs used in this study are trametinib (ChemieTek #CT-GSK212, IN, USA), 10058-F4 (MedChemExpress #HY-12702, NJ, USA), and abemaciclib (Selleckchem # S5716, TX, USA).

### Generation of trametinib resistant cell lines

To generate resistance, pancreatic cancer cells were incubated in growth media supplemented with 10 nM trametinib. Following an initial phase of growth arrest and cell death, a population emerged capable of proliferation, at which point the concentration of trametinib was increased to 20 nM. This process was repeated, with dose escalation in 20 nM to 50 nM increments until the concentration of trametinib was escalated up to 200 nM (188 and CFPAC-1) or 400 nM (395 and PANC-1). Cells were periodically transferred to trametinib-free media to allow for recovery and outgrowth. Generation of trametinib resistant cell lines took 4-6 months per cell line. Resistant cells were maintained on the highest concentration of trametinib to which they are resistant throughout the study.

### Western blot

Cell pellets were lysed on ice in RIPA (1% IGEPAL CA-630, 20mM Tris pH 8, 137mM NaCl, 10% glycerol, and 2mM EDTA) buffer with protease inhibitor cocktail (Milipore Sigma #11836170001, IN, USA) and phosphatase inhibitor cocktail (Thermo Fisher #A32957, IL, USA) Protein concentration was determined by Bio-Rad Protein Assay dye (Bio-Rad #5000006, CA, USA). Either an equal fraction of cells (100,000) or an equal amount of protein (20 μg or 15 μg) was resolved by SDS-PAGE on a 10% polyacrylamide gel, transferred to PVDF membrane (Millipore Sigma #IPVH00010), blocked in 5% fat-free milk, and probed with indicated antibodies. Primary antibodies used in this study are: P-DRP1 (S616) (CST #4494S, MA, USA), DRP1 (Abcam #184247, UK), Beta Actin (CST #8457), MFN1 (CST #14739), MFN2 (CST #9482), OPA1 (BD Biosciences #612606, NJ, USA), P-ELK-1 (S383) (CST #9181S), ELK-1 (CST #9182S), P-ERK (Thr 202/Tyr 204) (CST #9101S), ERK (CST #9102S), c-Myc (CST #5605), P-RB (S807/811) (CST #9308S), RB (CST #9309), CDK4 (CST #12790T), CDK6 (Santa Cruz #7961), CDK2 (SantaCruz #.6248) The secondary antibodies used in this study are: goat anti-rabbit HRP (Jackson ImmunoResearch #111-033-003, PA, USA), or goat anti-mouse HRP (Jackson ImmunoResearch #115-035-003). Western blots were developed using western bright ECL solution (advansta #R-03025-D10, CA, USA).

### MitoTracker Red staining and imaging

100,000 cells (188 cells) or 50,000 cells (PANC-1 cells) were seeded on glass slides (VWR #48366-227, PA, USA) in RPMI media. After overnight incubation, media was replaced with fresh media containing either DMSO or trametinib (200 nM). After 48 hrs. media was removed, cells were incubated in 50 nM MitoTracker Red CMXRos (invitrogen #M7512, MA, USA) for 30 minutes, washed in media containing the same concentration of drug for 1 hr., fixed for 15 min with 4% paraformaldehyde in media and washed 3 x 5 minutes with 1X PBS. At the last wash step, DAPI solution was added at a 1:20,000 dilution. After the final wash, slides were mounted immediately in Prolong Gold antifade reagent (CST #9071). A Zeiss LSM 900 with Airyscan2 confocal microscope was used for imaging mitochondrial network using a 63x objective.

### Mitochondrial morphology analysis

Mitochondrial structure analysis was performed using MitoHacker [27] through its web-based interface (mitogenie.com).

### Principal Component Analysis

To determine (in an unsupervised manner) what mitochondrial structural features contribute most significantly to the observed differences in mitochondrial morphology across different biological conditions, we performed principal component analysis (PCA). We first assembled a 2-dimensional data matrix with 108 columns (representing 108 mitochondrial features captured by MitoHacker) and 24 rows representing different experimental conditions, including 2 cell lines (188 and PANC-1) × 2 sensitivity conditions (resistant versus sensitive) × 2 treatment conditions (trametinib versus DMSO) × 3 independent replicates. The single-cell data for each mitochondrial feature were averaged to obtain a value for each experimental condition. The data was z-scored across all 24 experimental conditions. We then performed PCA using sklearn.decomposition.PCA from scikit-learn in Python, followed by visualization of PCA scores to illustrate how data are separated along PCs.

Because PC1 and PC2 separated treatments and cell lines, we identified the features that contributed most to these PCs by identifying PCA loadings with an absolute value > 0.1. To limit redundancy, we removed features which were highly correlated in mathematical nature. From these filtered features, we performed additional statistical analyses on those with the greatest positive loadings or smallest negative loadings, comparing them across biological conditions.

### CellTiterGlo (CTG) cell viability assay

1,000 cells were seeded in full RPMI in white-walled 96-well plates. After overnight incubation, media was removed, and cells were treated as indicated in the figure legend. The media (including indicated drug treatments) was changed every 3 days. At the end of the experiment, 50 μL of CTG reagent (Promega #G7572, WI, USA) diluted with 1xPBS (2:3) was added to each well and incubated for 10 min at room temperature prior to luminescence measurement (Perkin Elmer victor3).

### Plasmid Generation

pCW57-mDRP1-HygroR was generated from pCW57-mDRP1-BlastiR [28] by replacing the blasticidin resistance gene with the hygromycin resistance gene through infusion cloning. pLentiCRISPRv2 NeoR [28] was digested with the restriction enzyme BsmBI-v2 at 55⁰C (New England Biolabs #R0580, USA). Oligonucleotides containing control (CTR) or DRP1 target sequences were annealed by slow cooling and inserted into digested and purified pLentiCRISPRv2NeoR vectors with T4 DNA ligase (New England Biolabs #M0202T). A hygromycin-resistant version of pLenti-CMVTetR (Addgene #17492) was generated by replacing the blasticidin resistance gene with the hygromycin resistance gene with the through infusion cloning. To generate shRNA constructs, pSuper-retro-neo plasmid was generated by removing GFP coding sequence from pSuper-retro-gfp-neo (Oligoengenie #VEC-PRT-0005/0006). pSuper-retro-neo was digested with BglII and HindIII, and oligonucleotides encoding shCTR, shCDK4, shCDK6, shCDK2 or shc-Myc were annealed by slow cooling and inserted into digested and purified pSuper-retro-neo vector with T4 DNA ligase. sgTarget or shTarget sequences (From 5’ to 3’): sgCTR: GTATTACTGATATTGGTGGG sgDRP1: TCAGTGCTAGAAAGCCTGGT shCTR: CAACAAGATGAAGAGCACCAA shCDK4: GTTCTTCTGCAGTCCACATAT shCDK6: CATGAGATGTTCCTATCTTAA shCDK2: ACGGAGCTTGTTATCGCAAAT shc-Myc: CCCAAGGTAGTTATCCTTAAA

### Cell Line Transfection

To produce retrovirus, HEK-293T cells growing in DMEM were transfected with retroviral packing plasmid pCL-10A1 (Novus Biologicals #NBP2-29542) together with vector containing Moloney murine leukemia virus (MMLV) packaging signal (pCW57-mDRP1-HygroR or pSuper-reto-neo shCDK2, shCDK4, shCDK6 or sh c-Myc). To produce lentivirus HEK-293T cells were transfected with lentiviral packaging plasmids psPAX2 (Addgene #12260), pCMV-VSV-G (Addgene #8454), along with vector containing human immunodeficiency virus type 1 (HIV-1) packaging signal (pLentiCRISPRv2 sgCTR or sgDRP1, pLenti-CMV-TetR-HygroR). The next day, DMEM media was replenished. Media containing virus was collected after 24hrs and filtered with a 0.45μm syringe filter then added to target cells with polybrene (4 μg/ml) for 48hr. Cells were then selected by the addition of 500 μg/ml Geneticin (Gibco #10131035) or 500 μg/ml Hygromycin (Invitrogen #10687010). Uninfected control cells were used alongside to confirm selection.

### *In vitro* kinase assay

Recombinant his-tag CDK6/ GST-tag cyclinD (BPS-Bioscience #40097) was purchased. Recombinant GST-DRP-1^518–736^ and GST-DRP-1^518–736^ (SA mutant) was generated in our lab previously [29]. For *In vitro* kinase assay, 20 ng or 50 ng CDK6/cyclinD with 200 ng of purified GST-DRP-1^518–736^ or 50 ng CDK6/cyclinD with GST-DRP-1^518–736^ (SA mutant) were suspended in protein kinase buffer (20 mM Hepes-KOH buffer, pH 7.4, 15 mM EGTA, and 20 mM MgCl_2_) and incubated with 50 µM ATP, and 1 µM DTT for 30 min at 30°C. Samples were resolved by SDS-PAGE, and either immunoblotted for P-DRP1,and CDK6 or stained with Coomassie Brilliant Blue to visualize total recombinant DRP-1 protein.

### Orthotopic Xenograft

Mouse xenograft experiments were performed using a protocol that has been described previously [30,31]. Briefly, 1 million cells resuspended in 50 uL 50% full RPMI:50% Matrigel®Growth Factor Reduced Basement Membrane Matrix (356230 Corning, NY, USA) were injected into the pancreas of a 6-8 weeks old male athymic nude mouse. Mice were anesthetized using a ketamine/dexmedetomidine mix and acclimated on a heating pad before an incision was made on the left flank and the pancreas was exteriorized. Cells were then injected directly into the pancreas using a 29-gauge needle. The abdomen was closed in two layers before anesthetic reversal and ketoprofen analgesia were administered and the animal was allowed to recover on the heating pad until fully conscious. Treatment vehicle or trametinib (0.3 mg/kg) was given through oral gavage once a day for the duration of the study. After 4 weeks or 6 weeks, as indicated in figure legend, mice were euthanized, tumors excised and tumor weights were measured using a weighing scale. Animals were randomly assigned to groups and for the experiment in Fig. 6g, sample sizes were determined based on power calculations using the results from the pilot animal experiment in Fig. 6e. Investigators were not blinded to the treatment groups. One animal was excluded from analysis because it died of an unknown cause prior to tumor initiation.

### Quantification and Statistical Analysis

Image lab 3 software (Bio-Rad) was used to quantify immunoblots. GraphPad Prism 7 was used to produce graphical presentation of data and to perform statistical analysis. Data was normalized as indicated in figure legends.

## Results

### Trametinib resistant pancreatic cancer cells exhibit increased DRP1 S616 phosphorylation

To determine the role of DRP1-mediated mitochondrial fission in acquired resistance of PDAC cells to MAPK inhibition, we treated two patient-derived xenograft cell lines (188, 395) [16,26] and two additional patient-derived cell lines (PANC-1, CFPAC-1) [32–34] with escalating doses of trametinib over a period of four to six months. Using this approach, we were able to establish populations of cells for each of the lines that maintain viability in the presence of 200 nM-400 nM trametinib. Analysis of the half maximal cell viability (IC50) for trametinib for each of the established cell lines demonstrates a significantly increased IC50 for each of the resistant (R) cells compared to sensitive (S) cells (Fig.1a). As MAPK-dependent activation of DRP1 has been previously shown to promote PDAC cell proliferation [14,16,35] we next sought to determine whether DRP1 expression and activation are altered in sensitive and resistant cells. Consistent with previous work, MEK inhibition leads to a decrease in DRP1 S616 phosphorylation in all four sensitive cell lines. However, all four resistant cells exhibit significantly increased DRP1 S616 phosphorylation following trametinib treatment when compared to sensitive cell lines (Fig. 1b,1c). Interestingly, two of the resistant cell lines exhibit increases in total DRP1 levels (188R and PANC-1R) while the other two exhibit equivalent (CFPAC-1R) or decreased (395R) DRP1 levels, suggesting multiple mechanisms may contribute to the maintenance of DRP1 activity in resistant cell lines (Fig. 1b,1c). To explore whether alteration of mitochondrial fusion activity also contributes to trametinib resistance, we analyzed expression of mitochondrial fusion mediators MFN1, MFN2, and OPA1 in resistant cells but did not observe significant changes (Fig. 1d). The maintenance of DRP1 phosphorylation in resistant cells could potentially result from either reactivation of MEK-ERK signaling or from the activation of an alternative kinase. To distinguish these two possibilities, we measured the phosphorylation of ERK (an indicator of MEK activity) and the phosphorylation of ELK1 (an indicator of ERK activity). Notably, trametinib treatment significantly decreases P-ERK and P-ELK1 levels in all four resistant cell lines (Fig .1d) suggesting alternative mechanisms have arisen in these cells to maintain DRP1 phosphorylation.

**Fig. 1:**
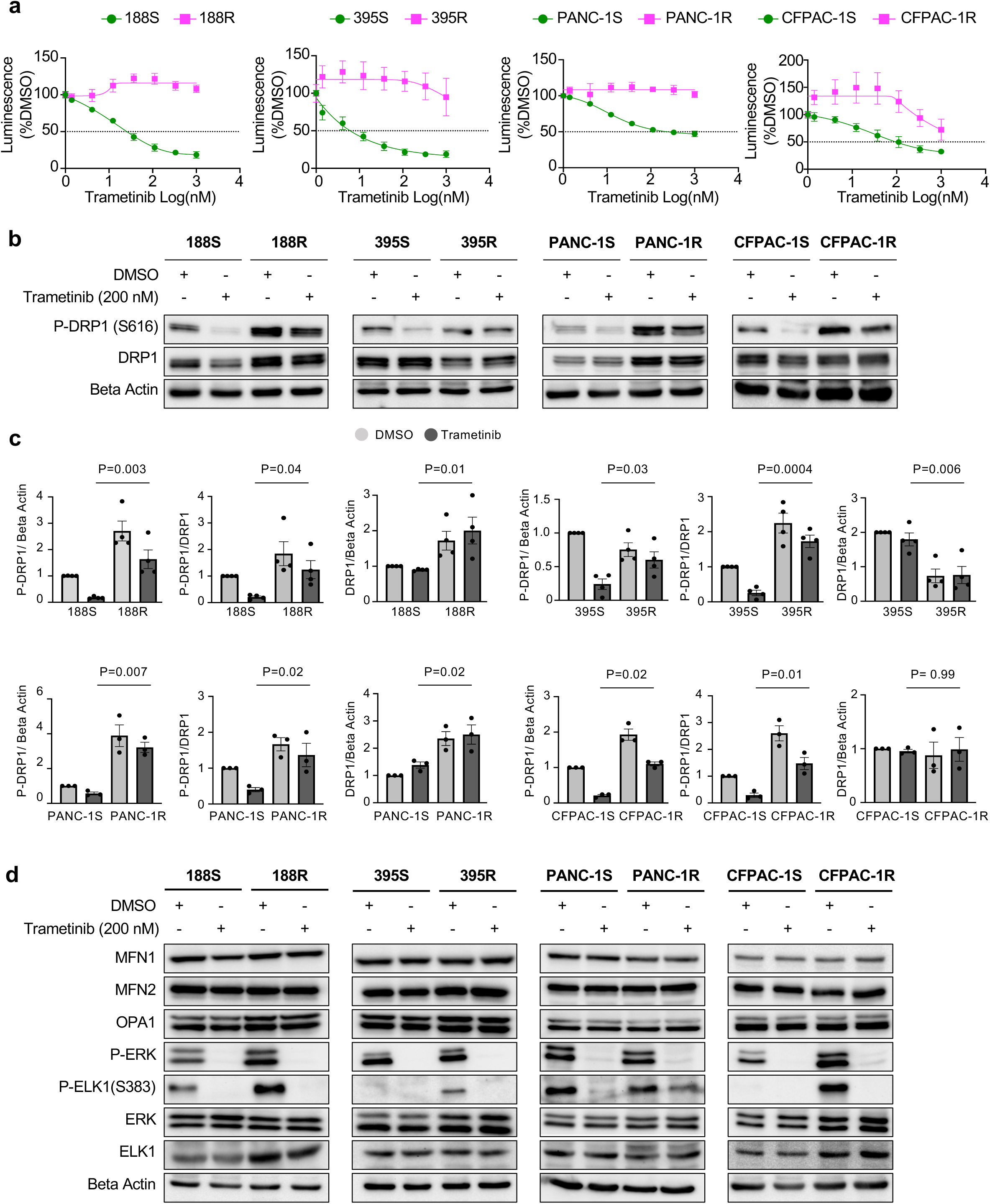
Trametinib resistant pancreatic cancer cells exhibit increased DRP1 S616 phosphorylation. a. Trametinib-sensitive and resistant patient-derived pancreatic cancer cell lines 188, 395, PANC-1 and CFPAC-1 were treated with DMSO or the indicated concentrations of trametinib for 5 days. Viable cells were quantified by CTG assay. N=6 wells per condition. Three technical replicates of each two independent experiments. b. Western blot analysis of P-DRP1 (S616) and total DRP1 from the indicated cells treated DMSO or trametinib (200 nM) for 24hrs. An equal fraction of cells (100,000) was loaded per lane. N=3 to 4 independent experiments. c. Quantification of the blots from b. Data was normalized to corresponding sensitive cells DMSO condition. Mean ± SD. Two-way ANOVA. d. Western blot analysis of the indicated proteins and post-translational modifications from the indicated cells treated DMSO or trametinib (200nM) for 24hrs. An equal fraction of cells (100,000) was loaded per lane. Representative blots from N=3 independent experiments.

### Trametinib-resistant pancreatic cancer cells exhibit distinct mitochondrial morphology compared to sensitive cells

One of the known functional outcomes of DRP1 S616 phosphorylation is mitochondrial fission, and previous studies have demonstrated that inhibition of MEK or ERK in PDAC, melanoma, breast, and colon cancer cells leads to significant increases in mitochondrial connectivity [14,15]. Hence, we next sought to analyze the mitochondrial morphology of resistant cell lines in response to trametinib using confocal microscopy in combination with analysis by MitoHacker [27], a computational pipeline that quantifies multiple metrics of mitochondrial structure and distribution from fluorescence images at the single cell level. MitoHacker analysis of images taken of two sets of resistant and sensitive cell lines (188 and PANC-1) treated with either vehicle control or 200 nM trametinib for 48 hours reveals significant differences in how mitochondrial structure is altered in resistant cells following trametinib treatment. The mitochondrial morphology of both trametinib-sensitive cell lines becomes visibly more connected and spread throughout the cytoplasm following trametinib treatment (Fig. 2a-b, Fig. S1a-b) consistent with previous literature. However, the trametinib-resistant lines appear visibly distinct from the sensitive lines in the presence of both DMSO and trametinib (Fig. 2a-b, Fig. S1a-b). To understand how trametinib resistance impacts mitochondrial morphology both at baseline and in response to treatment, we performed principal component analysis (PCA) on all mitochondrial features reported by MitoHacker, using the average values from over 100 individual cells per condition (Fig. 3a). By analyzing data from three independent replicates per condition, we were able to identify variations in mitochondrial features associated with biological conditions that exceeded experiment-to-experiment variability.

**Fig. 2:**
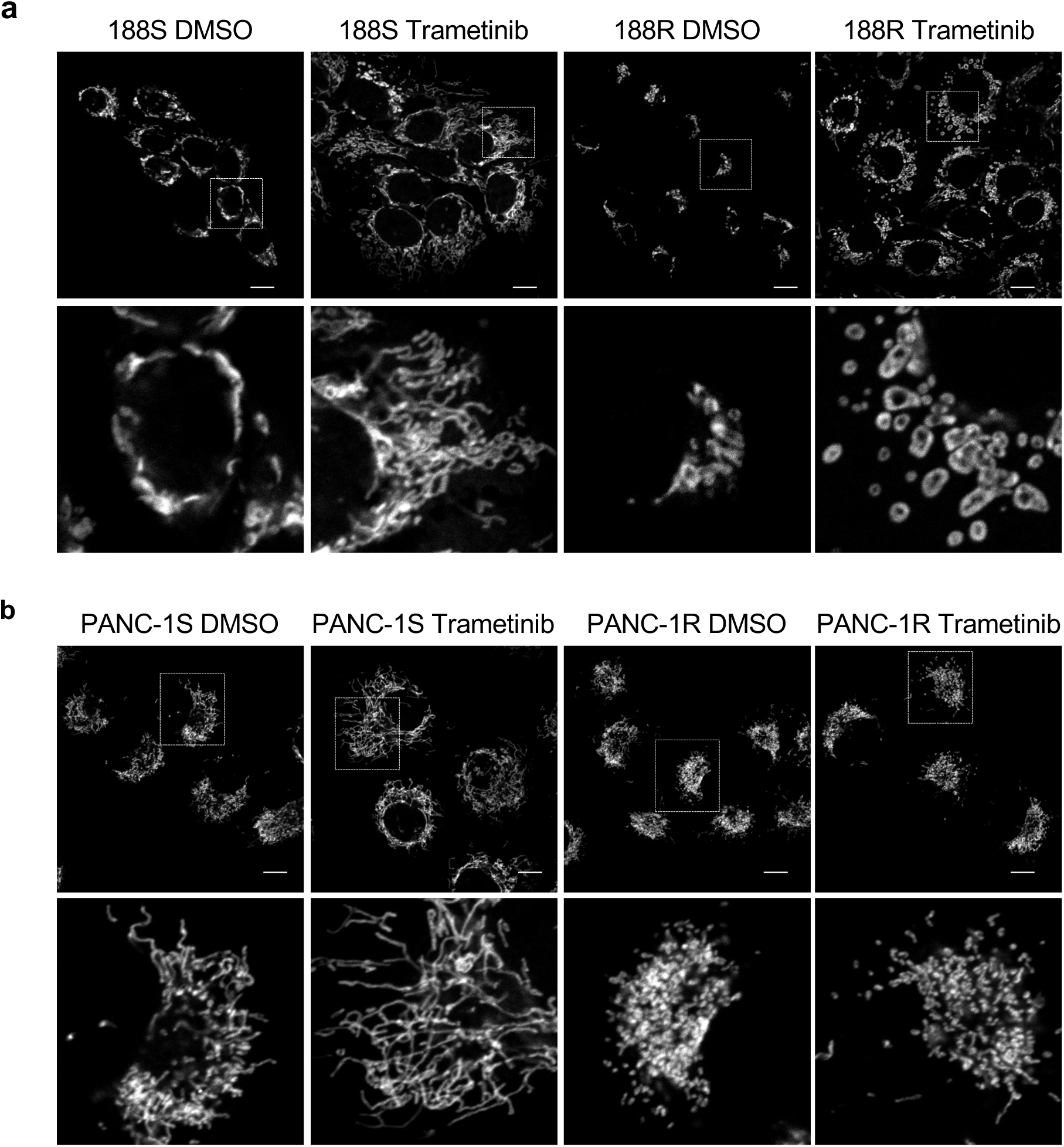
Trametinib has distinct effects on mitochondrial morphology in trametinib-sensitive and resistant pancreatic cancer cells. a. 188S and 188R cells were treated with either DMSO or trametinib (200 nM) for 48hr and stained with MitoTracker red to visualize the mitochondrial network. Representative images from N=3 independent experiments. 30 to 75 cells were analyzed per replicate for each condition. b. PANC-1S and PANC-1R cells were treated with either DMSO or trametinib (200 nM) for 48hr and stained with MitoTracker Red to visualize the mitochondrial network. Representative images from N=3 independent experiments. 30 to 75 cells were analyzed per replicate for each condition.

**Fig. 3:**
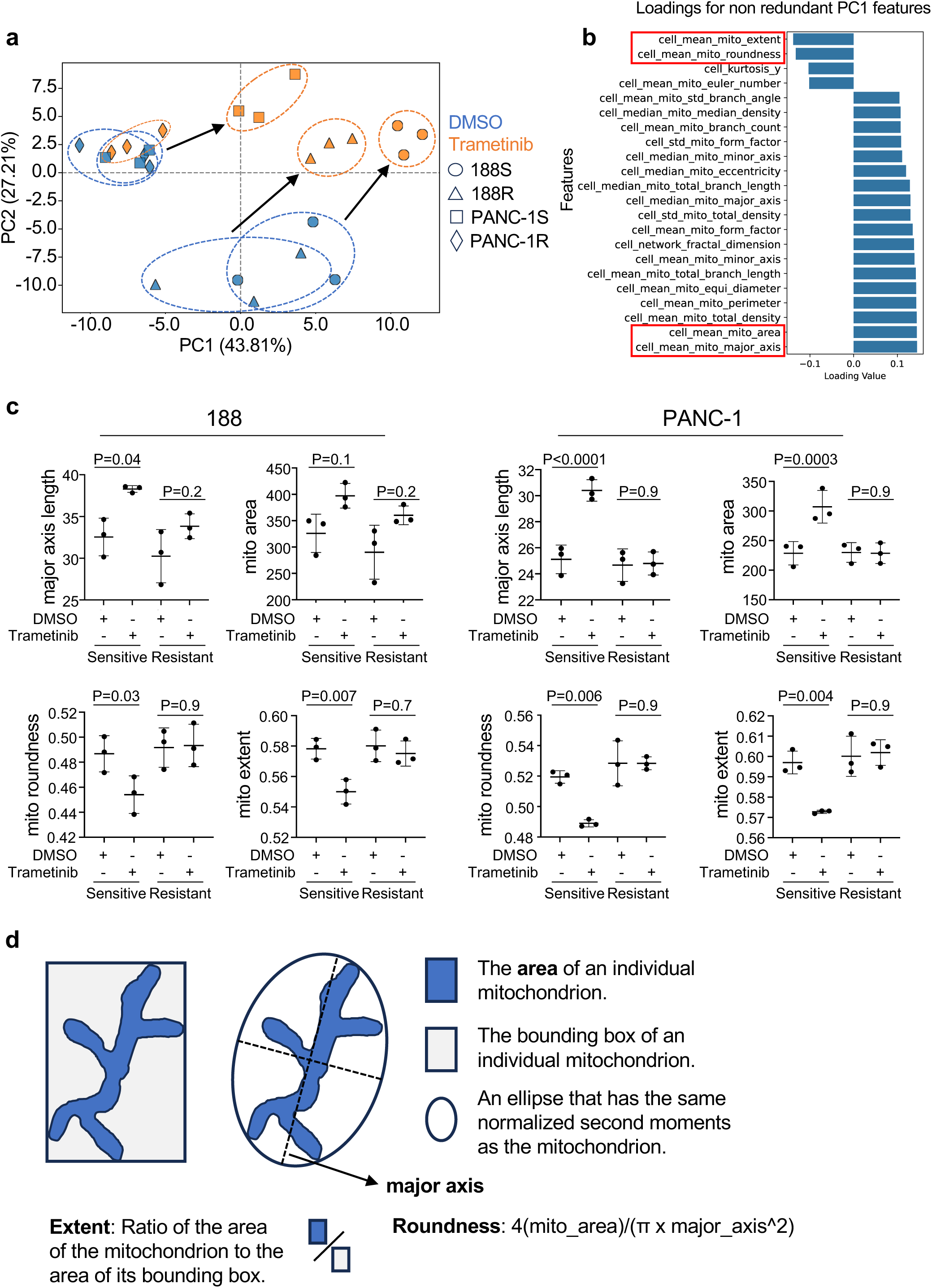
PCA of mitochondrial features reveals separation between trametinib-resistant and trametinib-sensitive cell lines and the effect of trametinib treatment. a. PCA scores across 24 conditions, including 2 cell lines (188 and PANC-1) × 2 sensitivity conditions (S: sensitive; R: resistant) × 2 treatment conditions (48 h treatment with trametinib at 200 nM, or DMSO), each tested in 3 replicates. Arrows highlight significant trametinib-induced changes in mitochondrial features. b. PC1 loadings of non-redundant mitochondrial features with the strongest influences on PC1. c. Representative graphs of the differentially regulated features following trametinib treatment. The single-cell data for each mitochondrial feature were averaged to obtain a value for each experimental condition. P values represent two-way ANOVA analysis of DMSO vs Trametinib for each cell line. d. Description of features highlighted in (c)

For 188 cells, sensitive (188S) and resistant (188R) populations were not separable under DMSO treatment (Fig. 3a). However, upon trametinib treatment, 188S and 188R cells became distinguishable from one another and from their DMSO-treated counterparts, with separation captured by mitochondrial features contributing to PC1 and PC2. Similarly, sensitive and resistant PANC-1 cells (PANC-1S and PANC-1R) were not separated under DMSO treatment but were distinguishable from 188 cells, consistent with the visible heterogeneity in mitochondrial morphology between cell lines. Notably, trametinib treatment led to a significant shift along PC1 and PC2 in PANC-1S cells, whereas PANC-1R cells showed little change relative to DMSO-treated controls. These data suggest that mitochondrial morphology in PANC-1R cells is relatively insensitive to trametinib treatment. To identify the structural features underlying these differences, we examined PCA loadings to determine which mitochondrial features contributed positively or negatively to PC1 and PC2 (Fig. 3b and Fig. S2a). Given that both resistant lines following trametinib treatment separated from their sensitive counterparts primarily along the PC1 axis, we focused on features that contributed most to PC1. As predicted by PCA, features associated with mitochondrial length and size, including major axis length (cell_mean_mito_major_axis), mitochondrial area (cell_mean_mito_area), and mitochondrial density (cell_mean_mito_total_density) ([27]) were consistently upregulated upon trametinib treatment in sensitive cells, but not in resistant cells (Fig. 3b-c). Conversely, mitochondrial extent (cell_mean_mito_extent) and roundness (cell_mean_mito_roundness), features associated with mitochondrial distribution (See definitions, Fig. 3d), were consistently downregulated upon trametinib treatment in sensitive cells but remain unchanged in resistant cells (Fig. 3b-c). Collectively, these data support a model in which trametinib resistance leads to changes in DRP1 activity which in turn leads to measurable and consistent changes in the structure of the mitochondrial network.

### c-Myc contributes to DRP1 S616 phosphorylation in trametinib resistant cells

Recent studies have identified that c-Myc is required for resistance to combined treatment with trametinib and hydroxychloroquine in pancreatic cancer [36] and that *Myc* copy number gain is a mechanism of resistance to the broad-spectrum RAS (ON) inhibitor RMC-7977 [37]. Consistent with this, we observe elevated levels of c-Myc in 188R, 395R and PANC-1R cells compared to their sensitive counterparts following trametinib treatment (Fig. 4a). Next, to determine if increased c-Myc activity contributes to the DRP1 S616 phosphorylation we observe in resistant cells, we treated 188R, 395R, and PANC-1R cells with the c-Myc inhibitor 10058-F4. Interestingly, inhibition of c-Myc leads to decreased levels of DRP1 S616 phosphorylation in 188R, 395R, and PANC-1R cells (Fig. 4b). Notably, c-Myc inhibition alone is sufficient to decrease DRP1 phosphorylation in all three cell lines. Furthermore, c-Myc inhibition has minimal impact on ERK phosphorylation in 2 of the 3 lines tested, consistent with our previous data indicating that an alternative kinase is responsible for maintaining DRP1 phosphorylation in trametinib-resistant cell lines. We further tested c-Myc inhibition in trametinib sensitive cells. Interestingly, inhibition of c-Myc in sensitive cells also leads to inhibition of DRP1 S616 phosphorylation suggesting that the c-Myc contribution to DRP1 S616 phosphorylation preexists in sensitive cells (Fig. S3a).

**Fig. 4:**
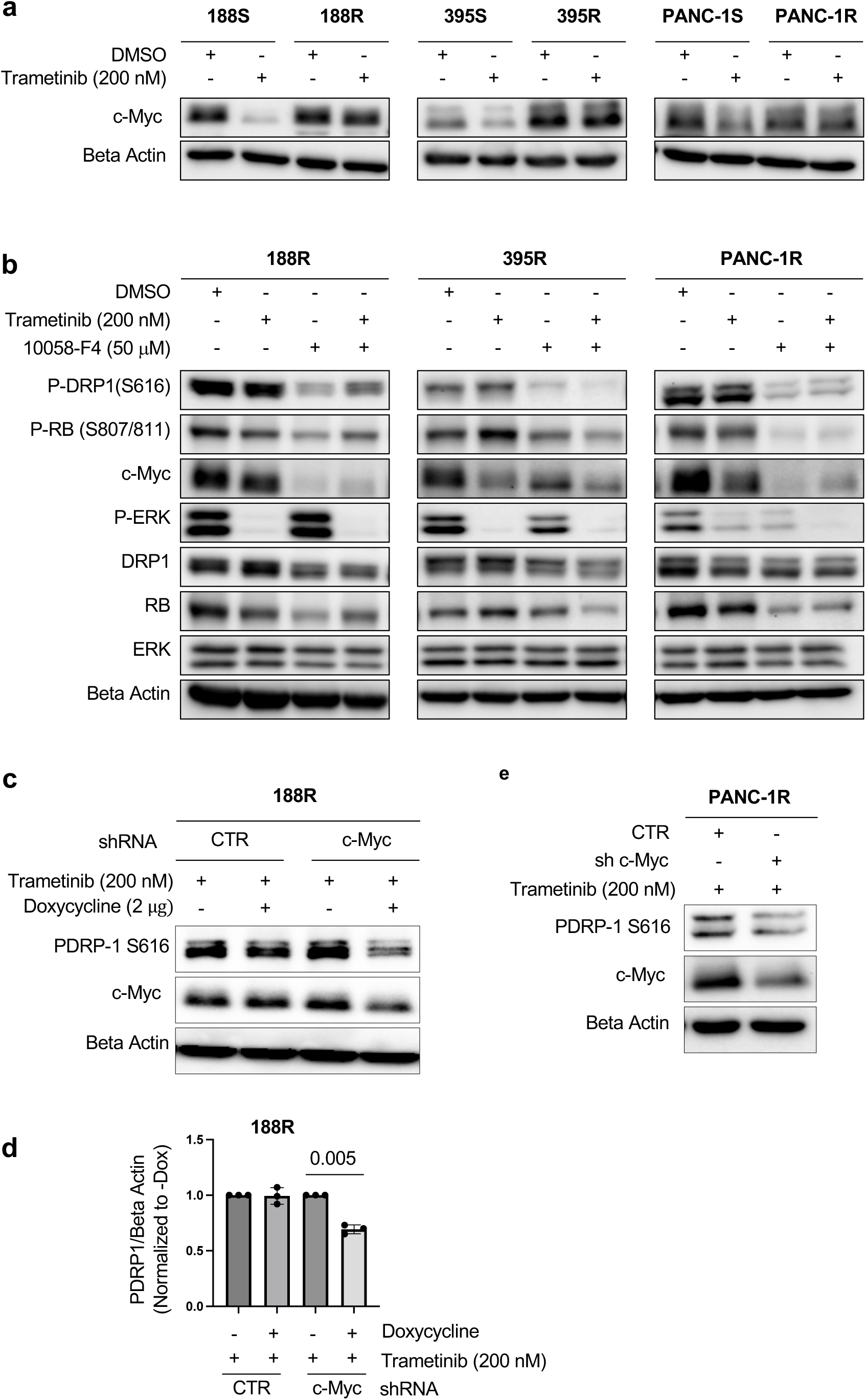
c-Myc contribute to DRP1 S616 phosphorylation in trametinib-resistant pancreatic cancer cells. a. Western blot analysis of c-Myc in 188, 395 and PANC-1 sensitive and resistant cells. An equal fraction of cells (100,000) was loaded per lane. N=3 independent experiments. b. Western blot analysis of the indicated proteins and post-translational modifications from trametinib-resistant 188, 395 and PANC-1 cells following 24hr treatment with either DMSO, trametinib (200 nM), 10058-F4 (50 μM), or a combination of trametinib (200 nM) and 10058-F4 (50 μM). 20 μg protein was loaded per lane. N=3 independent experiments. c. Western blot analysis of 188R cells expressing doxycycline inducible shRNA targeting a control sequence or c-Myc treated for 48 hrs with doxycycline to induce knockdown and trametinib (200 nM). N=3 independent experiments. d. Quantification of the blots from c. Welch’s t test. e. Western blot analysis of PANC-1R cells expressing shRNA targeting a control sequence or c-Myc were treated with trametinib (200 nM) for 24hrs.

To validate the impact of 10058-F4 on DRP1 phosphorylation is not an off-target effect, we engineered 188R cells to express doxycycline inducible shRNA targeting either c-Myc or a control sequence. Consistent with drug treated cells, we observe a significant reduction of DRP1 S616 phosphorylation following c-Myc knockdown (Fig. 4c-d). We also observed similar effect in PANC-1R cells where c-Myc knockdown leads to a decrease in DRP1 S616 phosphorylation (Fig. 4e). Collectively, these data demonstrate that c-Myc contributes to DRP1 S616 phosphorylation in MEKi resistant pancreatic cancer cells.

### CDK6 contributes to DRP1 S616 phosphorylation in trametinib resistant pancreatic cancer cells

c-Myc sustains tumor growth through engagement of multiple pro-proliferation pathways, including activation of the cyclin dependent kinases CDK4 and CDK6 [38–41]. DRP1 S616 has previously been identified as a target of several different kinases, including CDK1 and CDK5 [42,43] though whether other cyclin dependent kinases are able to phosphorylate this site has not been previously described. We tested whether inhibition of c-Myc also lead to inhibition of phosphorylation of RB at S807/811, a direct target of CDK4/6. We observed that inhibition of c-Myc in three of the resistant cells tested inhibited phosphorylation of RB (Fig. 4b). From this observation we aimed to test CDK4/6 inhibition in the resistant cell lines. Notably, treatment of trametinib-resistant 188R, 395R, and PANC-1R cells with abemaciclib, a CDK4/6 inhibitor, leads to inhibition of DRP1 S616 phosphorylation in all three resistant cells line tested, either alone or in combination with trametinib (Fig. 5a). Importantly, phosphorylation of RB (S807/811) is also inhibited by abemaciclib, confirming inhibition of CDK4/6. Additionally, MEK activity, as measured by ERK phosphorylation, is either fully or partially retained following abemaciclib treatment in all three cell lines, suggesting that the drug is not inhibiting DRP1 phosphorylation through indirect inhibition of MAPK signaling. Interestingly, abemaciclib also inhibits DRP1 S616 phosphorylation in trametinib-sensitive 188S and 395S cells, but not PANC-1S cells, suggesting that CDK4/6 may be the primary DRP1 S616 kinase in some pancreatic cancer cells (Fig. S3b). To further confirm the contribution of CDK4 and CDK6 to DRP1 S616 phosphorylation in resistant cells, we generated doxycycline-inducible 188R CDK4 and CDK6 shRNA knockdown cells as well as scramble control cell lines. While knockdown of either CDK4 or CDK6 decreases DRP1 S616 phosphorylation, we observe a greater decrease in DRP1 S616 phosphorylation upon knockdown of CDK6 (Fig. 5b, 5c). To determine whether CDK6 is capable of direct DRP1 phosphorylation, we incubated recombinant DRP1 c-terminus (aa518-736) with recombinant cyclinD1/CDK6 complex in the presence of ATP. We observe DRP1 S616 phosphorylation following incubation with cyclinD1/CDK6 (Fig. 5d). This phosphorylation is not detected on a S616A mutant. Collectively, these data are consistent with a model in which CDK6 directly phosphorylates DRP1 to maintain its phosphorylation in trametinib-resistant pancreatic cancer cells. Importantly, knockdown of another cell cycle kinase, CDK2, in 188R cells does not impact DRP1 S616 phosphorylation (Fig. 5e). These data demonstrate that cell cycle inhibition alone does not inhibit DRP1 S616 phosphorylation in trametinib resistant cells and further support the model that DRP1 S616 phosphorylation is a direct consequence of CDK6 activity.

**Fig. 5:**
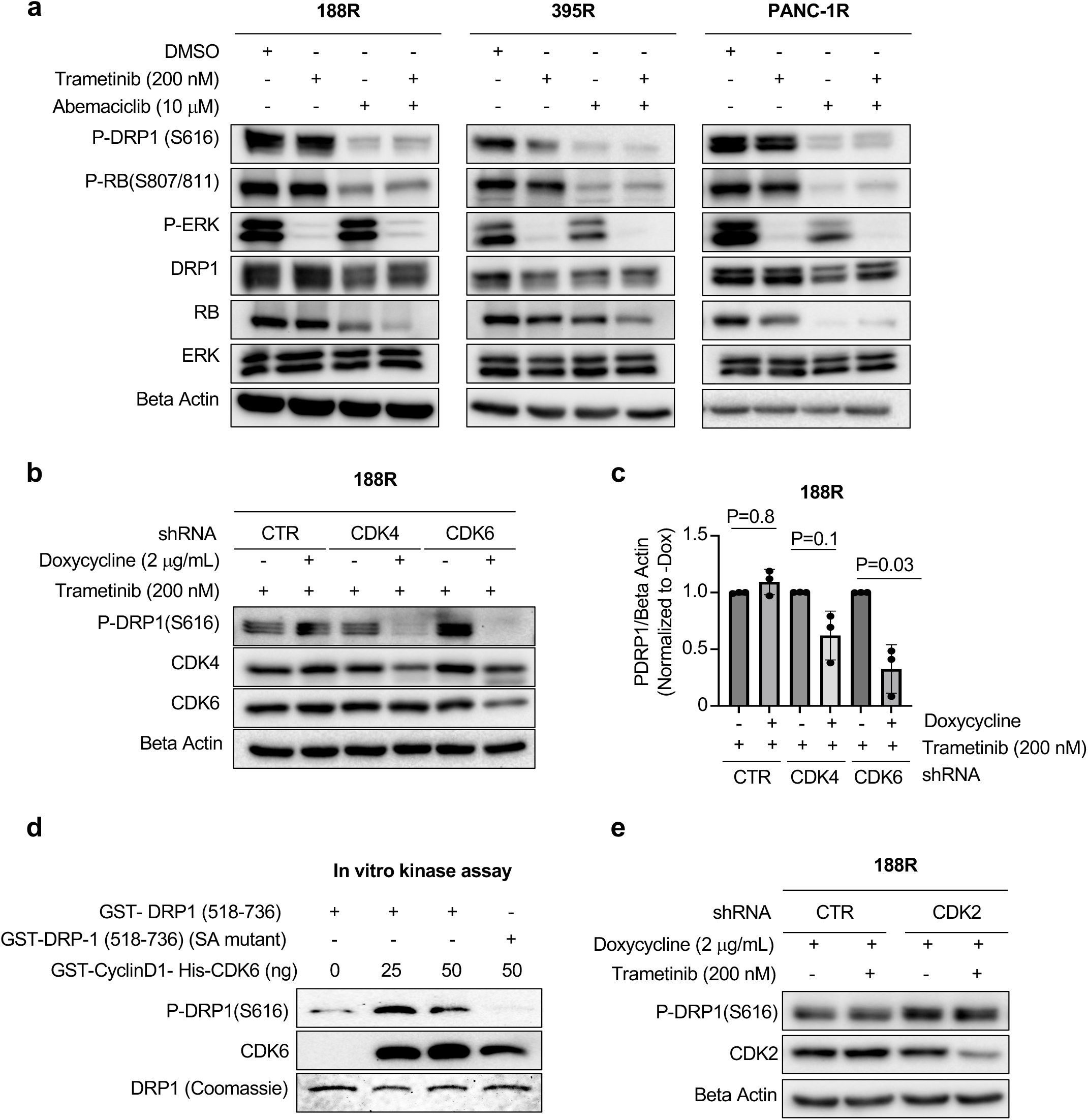
CDK6 promotes DRP1 S616 phosphorylation in trametinib-resistant pancreatic cancer cells. a. Western blot analysis of the indicated proteins and post-translational modifications from trametinib-resistant 188, 395 and PANC-1 cells following 24hr treatment with either DMSO, trametinib (200 nM), abemaciclib (10 μM) or combination of trametinib (200 nM) and abemaciclib (10 μM). 20 μg protein was loaded per lane. N=3 independent experiments. b. Western blot analysis of P-DRP1 (S616), CDK4 and CDK6 from trametinib-resistant 188 cells following doxycycline-induced expression of shRNAs targeting a control sequence, CDK4 or CDK6. 20 μg protein was loaded per lane. N=3 independent experiments. c. Quantification of the P-DRP1 levels from the experiment described in d. For each shRNA, P-DRP1 levels in doxycycline-treated cells were normalized to the corresponding untreated control. N=3 independent experiments. Welch’s t test was performed. d. In vitro kinase assay reaction including recombinant GST-DRP1 (518-736) (200 ng) and recombinant His-CDK6/GST-CyclinD1 (25 ng or 50 ng) in presence of ATP. GST-DRP1 (518-736) (SA mutant) was used as a negative control. Reactions were resolved by SDS–PAGE and subjected to either immunoblotting for P-DRP1 S616 and CDK6 or Coomassie staining to visualize recombinant DRP1 protein. N=3 independent experiments. e. Western blot analysis of P-DRP1 (S616) and CDK2 from trametinib-resistant 188 cells following doxycycline-induced expression of shRNA targeting a control sequence or CDK2. N=3 independent experiments. Welch’s t test was performed.

### Inhibition of DRP1 or CDK4/6 is able to resensitize or inhibit growth of the trametinib resistant cells

Given the increased DRP1 phosphorylation observed in trametinib resistant cells, we next sought to test the importance of DRP1 for the resistance phenotype. Because stable DRP1 knockout (DRP1KO) lead to metabolic adaptation of pancreatic cancer cells [16] we generated inducible DRP1KO trametinib-resistant cell lines by first stably expressing doxycycline-inducible mouse DRP1 (mDRP1) followed by expression of Cas9 and an sgRNA targeting either endogenous human DRP1 or a control sequence. Removal of doxycycline from the culture media for seven days results in near complete loss of DRP1 expression (Fig 6a). Interestingly, knockout of DRP1 causes growth arrest of three of the four cell lines tested (395R, PANC-1R, CFPAC-1R) in both the presence and absence of trametinib (Fig. 6b). These data are consistent with the requirement for DRP1 we have observed previously in pancreatic cancer cells [16]. Trametinib resistant 188R cells are insensitive to DRP1 knockout alone, but loss of DRP1 partially resensitizes these cells to trametinib, suggesting that the maintenance of DRP1 activity we observe is important for the resistant phenotype (Fig. 6b). Importantly, results from all four resistant cell lines suggests that inhibition of DRP1 is a potential approach to overcome therapeutic resistance to trametinib or other RAS-MAPK targeted therapies. Because inhibition of CDK4 and CDK6 lead to decreased DRP1 phosphorylation in trametinib-resistant cells, we next sought to test the effect of CDK4/6 inhibition on their growth. All three resistant cell lines tested were sensitive to abemaciclib (Fig. 6c). Consistent with the effects of DRP1 inhibition, abemaciclib treatment inhibits the growth of 188R, 395R and PANC-1R cells in both the presence and absence of trametinib (Fig. 6d). Collectively, these data are consistent with a model in which pancreatic cancer cells that have developed resistance to trametinib require CDK4/6 signaling to maintain their growth and the resistant phenotype requires reactivation of DRP1.

**Fig. 6:**
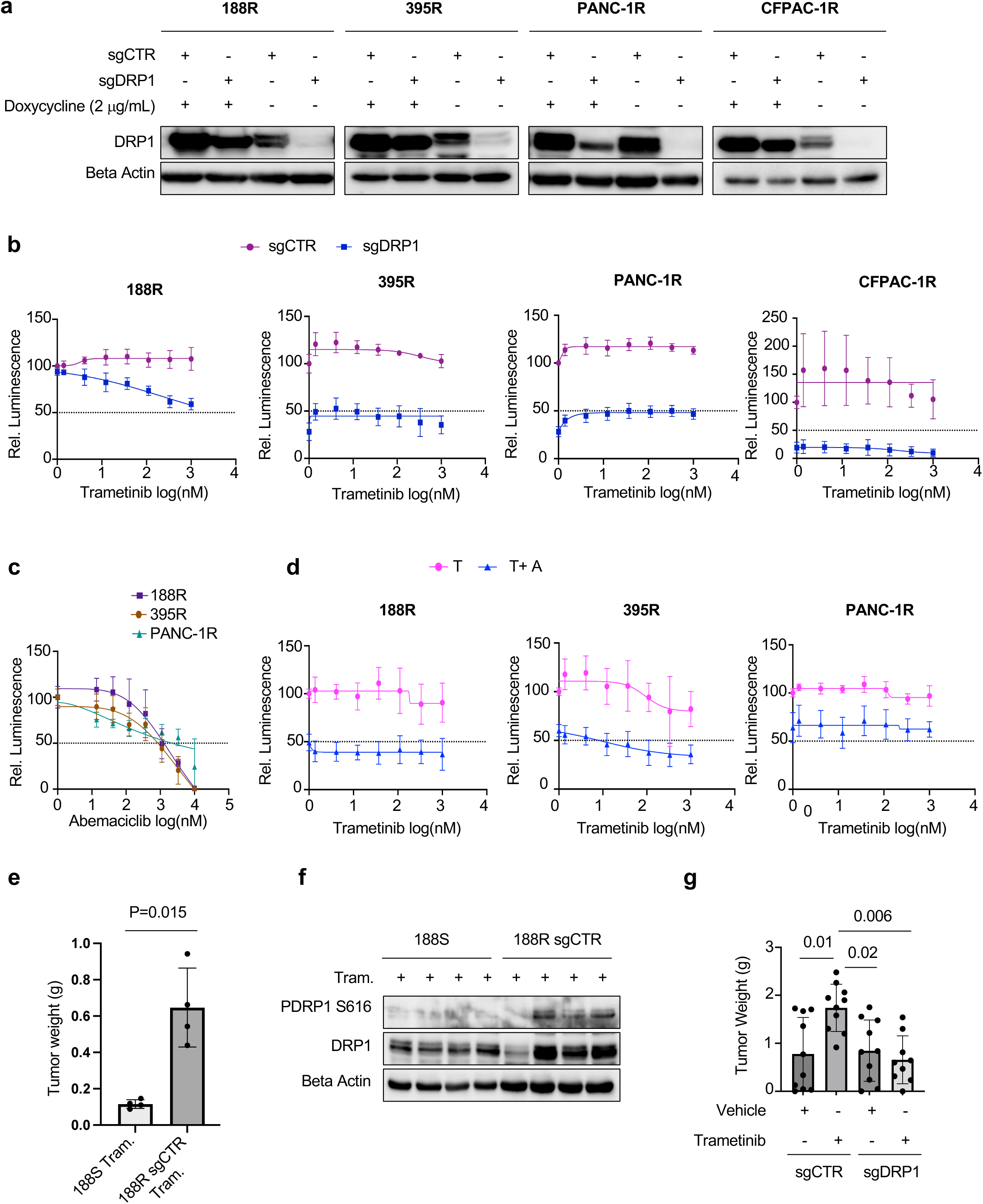
Inhibition of DRP1 and CDK4/6 in trametinib-resistant cells impacts growth and resistance phenotype. a. Western blot of DRP1 from trametinib-resistant 188, 395, PANC-1 and CFPAC-1 cells engineered to stably express a doxycycline-inducible mouse DRP1 and subsequently engineered to express CAS9 and CRISPR guides targeting either a control sequence or endogenous human DRP1. Cells were cultured in the presence or absence of doxycycline for 7 days prior to analysis. b. Inducible DRP1 knockout, trametinib-resistant 188, 395, PANC-1 and CFPAC-1 cells described in (a) were cultured in the absence of doxycycline for 7 days then treated with DMSO or the indicated concentrations of trametinib for 6 days (PANC-1R) or 9 days (188R, 395R, and CFPAC-1R). Viable cells were quantified by CTG assay. N=6 wells per condition. Mean±SD. Three technical replicates with two independent experiments c. Trametinib-resistant 188, 395 and PANC-1 cells were treated with DMSO or a range of abemaciclib doses for 5 days. Viable cells were quantified by CTG assay. N=4 wells per condition. Three independent experiments. Mean±SD. d. Trametinib-resistant 188, 395 and PANC-1 cells were treated with DMSO or a range of trametinib doses either alone, or in combination with 1 μM abemaciclib for 5 days. Viable cells were quantified by CTG assay. N=4 wells per condition. Three independent experiments. Mean±SD. e. 188S and 188RsgCTR cells were injected into the pancreata of nude mice (10^6^ cells per mouse) and mice were treated with trametinib (0.3 mg/kg) orally once a day for 4 weeks at which point tumors were harvested and weighed. Unpaired t test, Mean±SD f. Tumors harvested from mice described in (e) were lysed in RIPA and the levels of P-DRP1 (S616) and total DRP1 were analyzed by Western blot. g. 188R sgCTR and 188R sgDRP1 cells were cultured in the absence of doxycycline for 7 days to eliminate expression of mDRP1 and then 10^6^ cells were injected cells into the pancreata of nude mice. Mice were treated with either vehicle or trametinib (0.3mg/kg) orally once a day for six weeks and tumors were harvested and weighed. ANOVA. Mean±SD

Next, to test if the upregulation of DRP1 S616 phosphorylation we observe *in vitro* is also maintained *in vivo*, we orthotopically injected trametinib-sensitive 188S cells or trametinib-resistant 188RsgCTR cells (removed from doxycycline for 7 days) into the pancreas of nude mice and treated mice with trametinib once a day by oral gavage. Importantly, trametinib-resistant cells maintained their resistance *in vivo*, as mice injected with these cells exhibited a significant increase in tumor burden compared with mice injected with sensitive cells (Fig. 6e). Furthermore, we observed that tumors from 3 of the 4 mice injected with trametinib-resistant tumor cells, but none of the tumors harvested from mice injected with sensitive cells exhibited elevated DRP1 S616 phosphorylation, consistent with our *in vitro* observations (Fig. 6f). Because trametinib resistant cells maintained both their resistance and DRP1 phosphorylation *in vivo*, we next sought to test whether the *in vivo* resistance phenotype was dependent on DRP1. To test this, we removed doxycycline from 188R sgCTR and 188R sgDRP1 cells for seven days to remove expression of mDRP1 then orthotopically injected cells into the pancreas of nude mice. We then treated mice with either vehicle or trametinib once a day by oral gavage. Notably, mice injected with 188R sgCTR cells and treated with trametinib exhibit increased tumor growth compared with vehicle treated mice, suggesting resistant cells exhibit trametinib addiction in vivo, consistent with prior literature [44]. Importantly, mice injected with 188R sgDRP1 cells and treated with trametinib exhibit significantly lower tumor burden than trametinib treated mice injected with sgCTR cells (Fig. 6g) demonstrating a functional importance for DRP1 expression *in vivo*. Collectively, these data are consistent with a model in which maintenance of DRP1 S616 phosphorylation plays a functional role in the maintenance of resistance to trametinib treatment in pancreatic cancer.

## Discussion

DRP1-mediated mitochondrial fission has been shown to be a key biological process that contributes to the growth of several different RAS-MAPK driven malignancies. In support of this, we now demonstrate that sustained DRP1 phosphorylation is a feature of pancreatic cancer cells that have developed resistance to RAS-MAPK targeted therapy and that resistant PDAC cells are vulnerable to DRP1 inhibition. Furthermore, we have shown that the c-Myc-CDK4/6 signaling axis is activated to maintain DRP1 activation in these cells. Collectively, this work suggests that targeting mitochondrial fission may be an attractive option for combating resistance to RAS-MAPK inhibition in pancreatic cancer and other RAS-driven malignancies.

A number of recent studies have identified pro tumorigenic roles for DRP1 in a variety of different tumor types [16–19]. Our observations that DRP1 phosphorylation is maintained in trametinib-resistant tumor cells and that these cells are sensitive to DRP1 inhibition are consistent with a general dependency on DRP1-dependent mitochondrial fission to maintain RAS-MAPK driven tumor growth. The exact mechanisms through which DRP1 activity promotes survival and proliferation in resistant cells is not explored here, but there are several potential pro-tumorigenic mechanisms that have been previously described. For example, we previously identified that DRP1 is required for RAS-induced glycolytic shift at least in part through facilitating increased transcription of the glycolytic enzyme hexokinase II [16]. In addition, DRP1 promotes mitophagy, a selective form of autophagy through which cells remove damaged mitochondria and salvage nutrients under stress, which has been shown to support tumor growth [45,46].

In agreement with a pro-fission role for DRP1 S616 phosphorylation we observe significant restructuring of the mitochondrial network in sensitive cells following trametinib treatment (Fig. 2 & 3) consistent with observations from previous analyses of pancreatic cancer cell lines treated with inhibitors of RAF, MEK or ERK [14,47]. Notably, analysis of mitochondrial structure in cell lines that have acquired resistance to MEK inhibition reveals distinct morphological responses to drug treatment, confirming the functional consequences of the sustained DRP1 phosphorylation we observe in the presence of trametinib. MitoHacker analysis of the mitochondrial features underlying the differences in mitochondrial structure demonstrates that resistant cells treated with trametinib have decreased length and area, consistent with a maintenance of fission activity. Phosphorylation of DRP1 at S616 has been associated with increased mitochondrial fission activity in a number of contexts, though the mechanism through which it promotes fission is not fully understood. Indeed, one recent study suggests that for certain isoforms and in certain contexts, this phosphorylation may negatively regulate DRP1 GTPase activity [48]. It should be noted that both resistant lines tested exhibit heterogenous mitochondrial morphology indicating that additional genetic and environmental factors are likely contributing to the overall structure of the mitochondrial network in pancreatic cancer cells.

We have identified mitochondrial structure as an attractive target to combat resistance to RAS-MAPK targeted therapies in pancreatic cancer. We observed that targeting DRP1 in resistant cells can either resensitize them to trametinib or directly inhibit cell growth. These results agree with previous observations that knockdown of DRP1 can inhibit pancreatic tumor growth *in vitro* and *in vivo* [14,16], though further investigations will be required to identify the factors that distinguish cells in which DRP1 inhibition alone inhibits growth versus those in which additional inhibition of MEK is required.

Results from our present study support a model sustained DRP1 activity alters mitochondrial structure and contributes to MEKi resistance. It is important to note that additional evidence will be required to determine whether mitochondrial structure itself or some function of DRP1 function independent of mitochondrial structure drives the resistant phenotype. Though this question is not easily addressable using classical genetic approaches, future work using high-content imaging and unbiased data-driven approaches should allow us to more directly evaluate the impact of complex heterogenous mitochondrial structure on cellular phenotypes independent of genetical or pharmacological interventions.

In this study, we provide evidence that DRP1 S616 phosphorylation in trametinib resistant cell lines is dependent on CDK4 and CDK6. While these kinases have not been previously implicated in direct phosphorylation of this site, it should be noted that other proline-directed cyclin dependent kinases, including CDK1 and CDK5 have been reported to target this site directly [42,43] and the sequence surrounding S616 aligns well with the consensus sites identified for CDK4 and CDK6 [49]. CDK4 and CDK6 share 70% sequence similarity [50] and most of the clinically developed drugs inhibit both kinases. Our genetic experiments suggest DRP1 phosphorylation in resistant cells is more dependent on CDK6 than CDK4, though the data suggests both kinases contribute to the phosphorylation in resistant cells. Notably, *CDKN2A*, the gene that encodes the CDK4/6 inhibitor p16 is frequently mutated in PDAC (46-60%) [51], suggesting the intriguing possibility that CDK4/6 may contribute to DRP1 activation even prior to the development of resistance. Consistent with this idea, we observed that both MEK inhibition and CDK4/6 inhibition were able to inhibit DRP1 S616 phosphorylation in two of the sensitive lines tested (188 and 395). These results suggest CDK4/6 may act as a direct DRP1 kinase in a subset of sensitive PDAC cells and that PDAC cells have multiple ways of activating DRP1. In a preclinical study, the combination of MEK inhibition and CDK4/6 inhibition was found to promote an increased survival advantage compared to monotherapy [52]. In another study, the combination of ERK inhibition and CDK4/6 inhibition was found to suppress PDAC cell line and organoid growth [53]. While a trametinib and CDK4/6i combination therapy proved ineffective in a very small cohort of metastatic PDAC patients [54], there is evidence that a personalized targeted therapy approach that matches therapy to specific genomic alterations might provide therapeutic benefits [55].

Our study has identified that c-Myc protein levels are elevated in resistant cells and contribute to DRP1 reactivation. MYC has emerged as a common mediator of resistance to RAS-MAPK pathway inhibition. For example, in BRAFi resistant melanoma, multiple resistance pathways, including reactivation of ERK, activation of PI3K, or activation of Notch1, converged upon a rebound of MYC [56]. This study further identified glycolysis as one of the MYC-dependent metabolic vulnerabilities partly due to the MYC dependent upregulation of hexokinase II. Given the previously identified role of DRP1 in maintaining RAS-driven hexokinase II expression, it is intriguing to speculate that c-Myc mediated reactivation of DRP1 contributes to glycolytic flux in these cells. In addition, c-Myc was previously reported to directly bind to the promoter region of DRP1 and contribute to transcriptional regulation of DRP1 [57]. The increased DRP1 protein expression we observe in trametinib resistant PDAC lines may thus represent an additional mechanism through which c-Myc maintains DRP1 activity in cells that have acquired resistance to RAS-MAPK targeted therapies, including the newly developed inhibitors that can directly target RAS, and which have opened a new avenue to combat RAS signaling in cancer. Notably, *c-Myc* copy number gain was found to be a mechanism of resistance to recently developed RAS-ON inhibitors, suggesting c-Myc-mediated DRP1 activation may be a shared mechanism of resistance to drugs that target this pathway [37].

Our finding that both c-Myc and CDK6 contribute to DRP1 S616 phosphorylation in resistant cells, and that c-Myc inhibition reduces both DRP1 phosphorylation and RB phosphorylation, suggests that increased c-Myc activity is upstream of CDK6 activity. However, previous studies make clear that c-Myc and CDK4/6 can regulate each other. For example, c-Myc null cells exhibit decreased CDK4/6 activity [58] whereas CDK4/6 knockdown cells leads to upregulation of gene signatures associated with Myc [59]. It is possible that these two factors exhibit reciprocal regulation in resistant PDAC cells and future experiments will be required to fully deconvolute their respective roles.

Analysis of tumor growth in vivo confirms a functional role for DRP1 in trametinib-resistant pancreatic tumors as DRP1 knockout significantly lowers tumor burden compared to sgCTR cells in mice treated with trametinib. Interestingly, sgCTR resistant cells grow significantly better in mice treated with trametinib than mice treated with vehicle, mirroring the “addiction” to RAS-targeted therapy that has been observed in other studies [44,60].

In conclusion, we have identified that a c-Myc-CDK6-DRP1 signaling axis plays a role in MEK inhibitor resistant pancreatic cancer cells, highlighting the importance of mitochondrial dynamics in pancreatic tumor growth and suggesting new avenues for therapeutic intervention.

## Acknowledgements

We thank the UVA department of Microbiology, Immunology and Cancer Biology for seed funding and microscopy support. We thank Dane T. Sessions, Christopher Prevost, Daniel Phipps, Hannah Coalson and Vicki Remley for intellectual support and suggestions. Graphical abstract of the study was generated by BioRender. The abstract of this study was published at, Cell Symposia (Multifaceted mitochondria) 2024 Sitges, Spain and AACR annual meeting 2025, Chicago, IL.

## Authors’ contributions

S.S. and D.F.K. conceptualized the project. S.S. generated reagents and performed the majority of the experiments. J.A.K. contributed to reagent generation and *in vitro* experiments. S.J.A. contributed to in vivo experiments. T.W.B. contributed to data interpretation and manuscript writing. E.B.L. and M.F.S. contributed to computational analysis on mitochondrial morphology. S.S., J.A.K. and D.F.K. interpreted the data. S.S. and D.F.K. wrote the manuscript.

## Ethics approval and consent to participate

Collection of human PDAC specimens was performed with approval of the Institutional Review Board at the University of Virginia in coordination with the Biorepository and Tissue Research Facility. All patients provided written consent for participation and no patients received neoadjuvant therapy. This study was carried out in strict accordance with the recommendations in the Guide for the Care and Use of Laboratory Animals of the National Institutes of Health. The protocol was approved by the Animal Care and Use Committee of the University of Virginia (PHS Assurance #A3245-01).

## Consent for publication

### Data availability

Further information about the data is available upon request to corresponding author David F. Kashatus

### Competing interests

The authors declare no competing interest

### Funding information

This work was supported by pilot funding from the UVA cancer center (to D.F.K. and S.S.), a Parsons-Weber-Parsons scholarship (S.S.), The Lustgarten Foundation (Innovation and Collaboration Program, to D.F.K. and T.W.B.) and the N.C.I. (U54CA274499-01-9941, to D.K.), a Farrow Fellowship from the UVA Comprehensive Cancer Center (S.S.), and the NCI Cancer Center Support Grant (P30 CA44579).

## Figure Legends

**Fig. S1.**
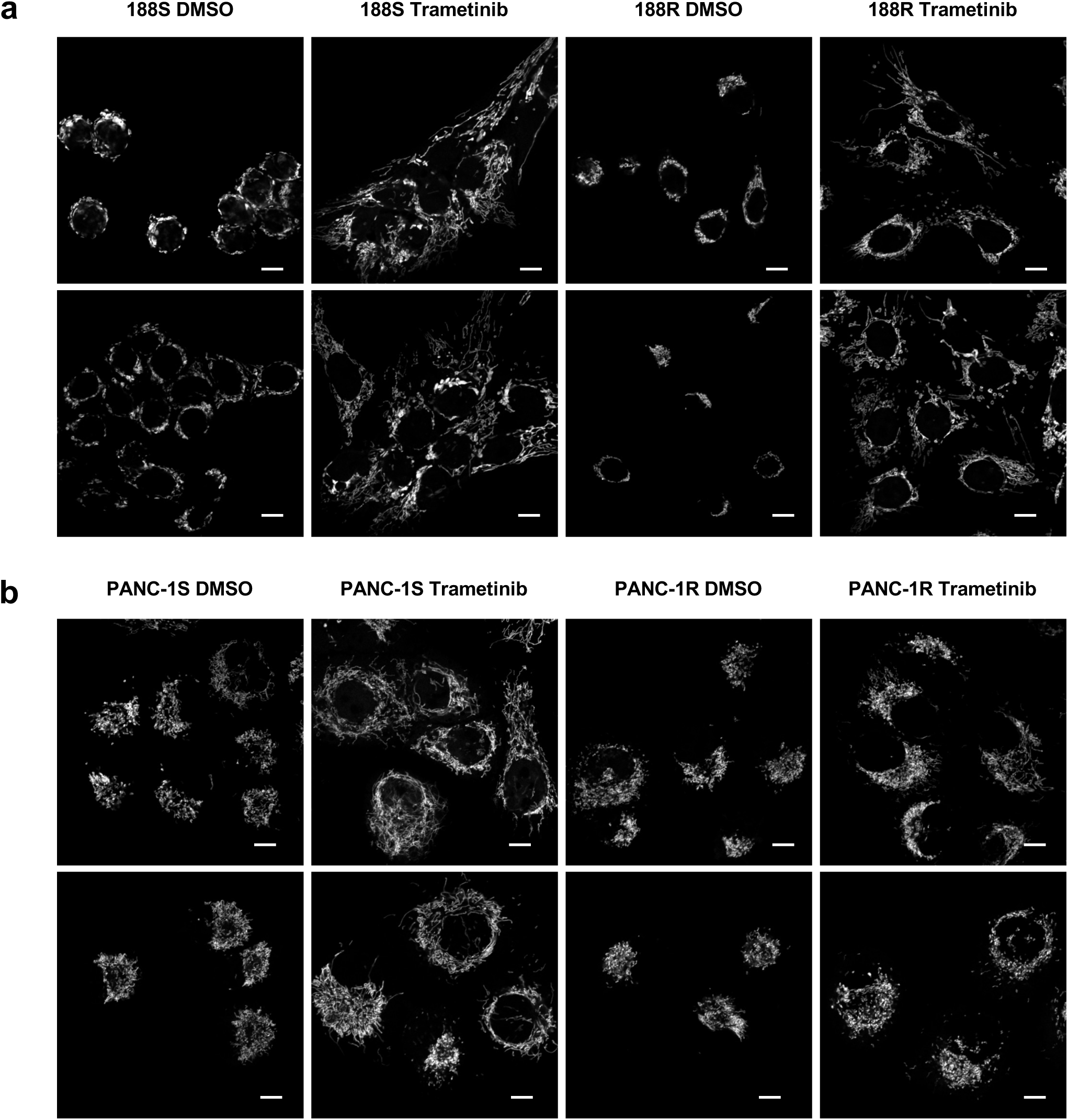
Trametinib has distinct effects on mitochondrial morphology in trametinib-sensitive and resistant pancreatic cancer cells. 188S and 188R cells (a) or PANC-1S and PANC-1R cells (b) were treated with either DMSO or trametinib (200 nM) for 48hr and stained with MitoTracker red to visualize the mitochondrial network. Representative images from N=3 independent experiments. 30 to 75 cells were analyzed per replicate for each condition.

**Fig. S2.**
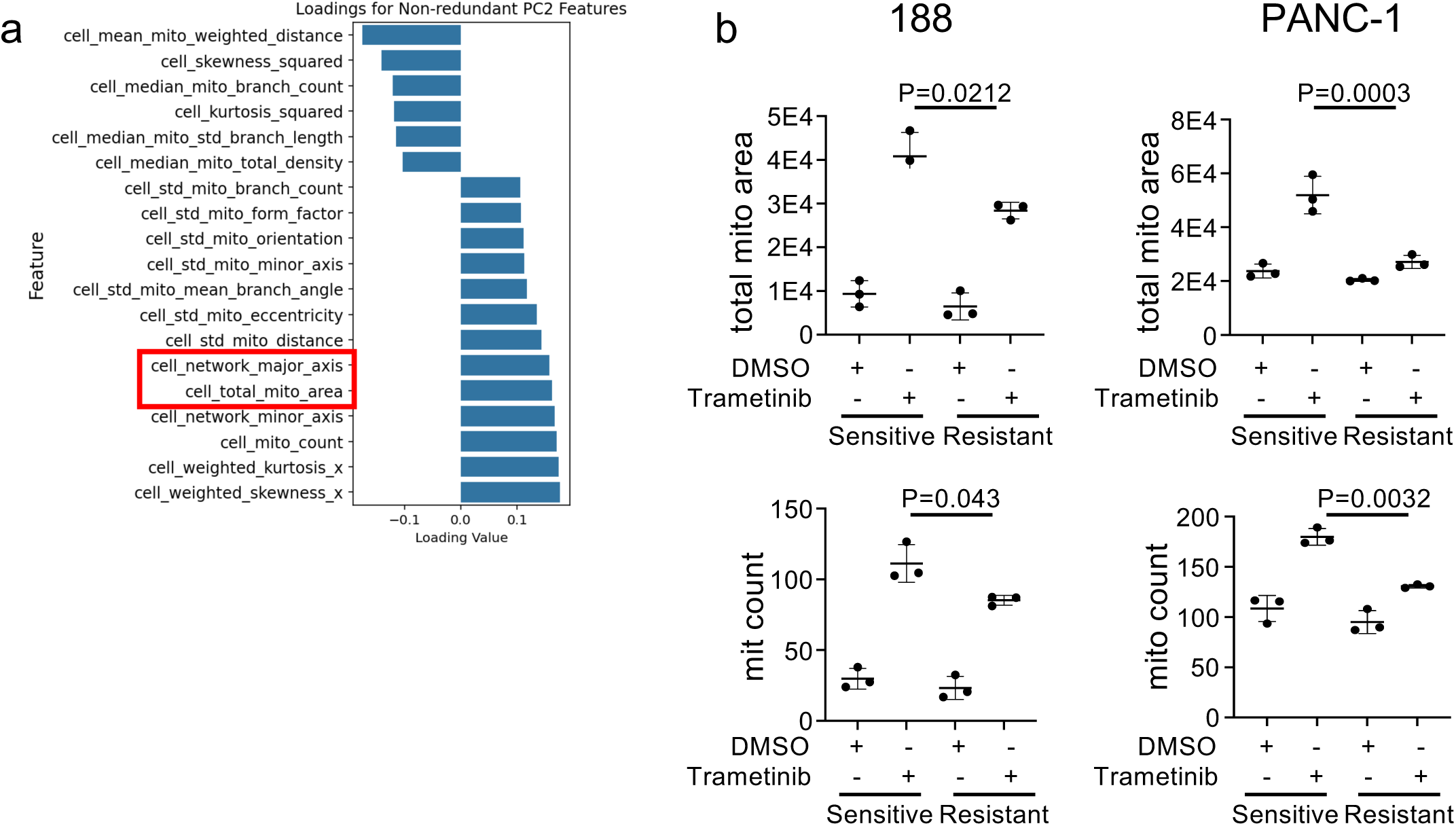
PCA of mitochondrial features reveals separation between trametinib-resistant and trametinib-sensitive cell lines. PC2 loadings (a) of non-redundant mitochondrial features with the strongest influences on PC2 from the principal component analysis in Figure 3. (b) Representative graphs of differentially regulated features following trametinib treatment. Single-cell data for each mitochondrial feature were averaged to obtain a value for each experimental condition. P values represent two-way ANOVA analysis of sensitive vs resistant cell lines treated with trametinib.

**Fig. S3.**
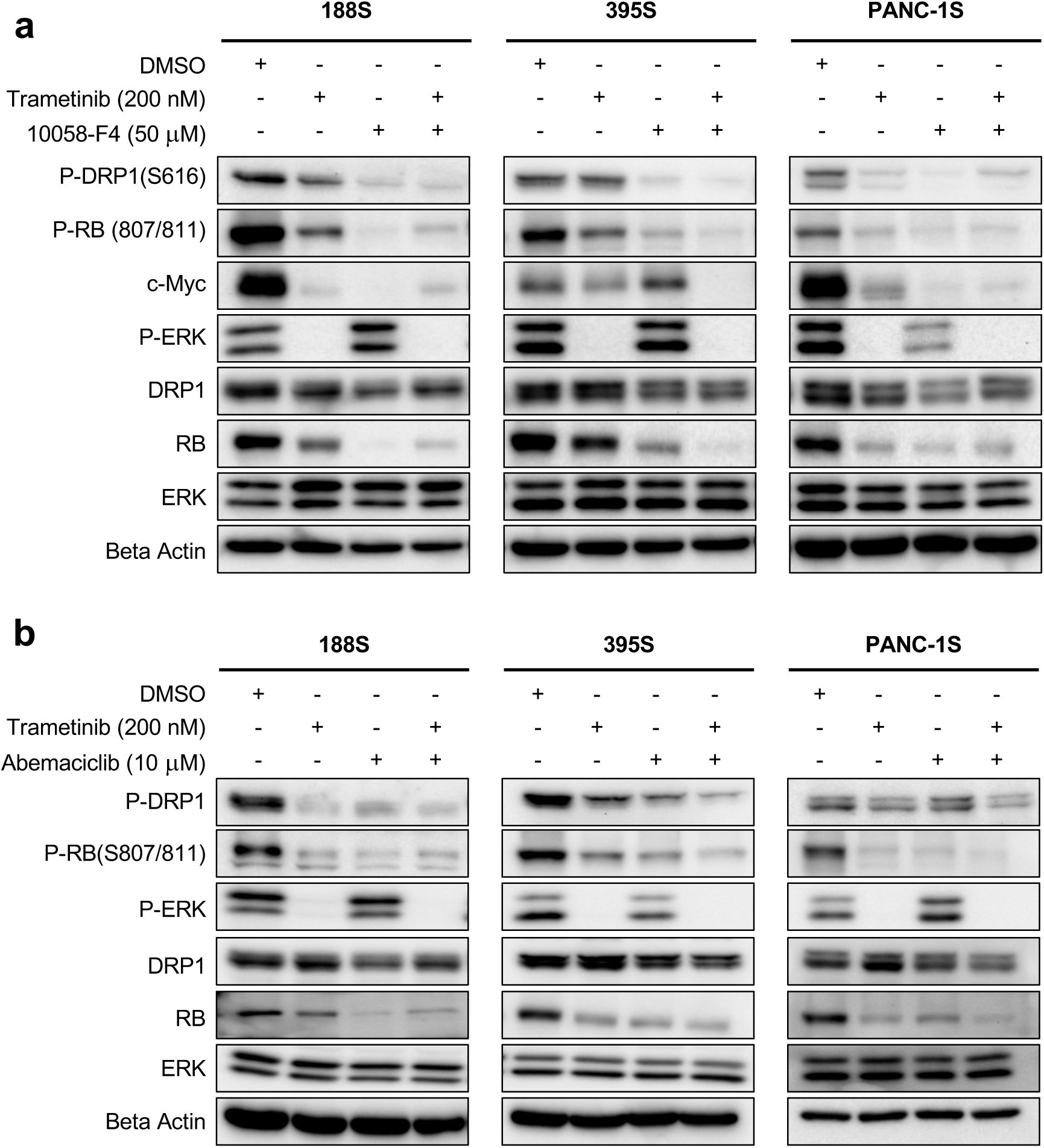
c-Myc and CDK4/6 contribute to the DRP1 S616 phosphorylation in trametinib-sensitive pancreatic cancer cells. Western blot analysis of the indicated proteins and post-translational modifications from trametinib-resistant 188, 395 and PANC-1 cells following 24hr treatment with: (a) DMSO, trametinib (200 nM), 10058-F4 (50 μM), or a combination of trametinib (200 nM) and 10058-F4 (50 μM); or (b) DMSO, trametinib (200 nM), abemaciclib (10 μM) or combination of trametinib (200 nM) and abemaciclib (10 μM). 20 μg protein was loaded per lane. N=2 independent experiments.

**Fig. S4.**
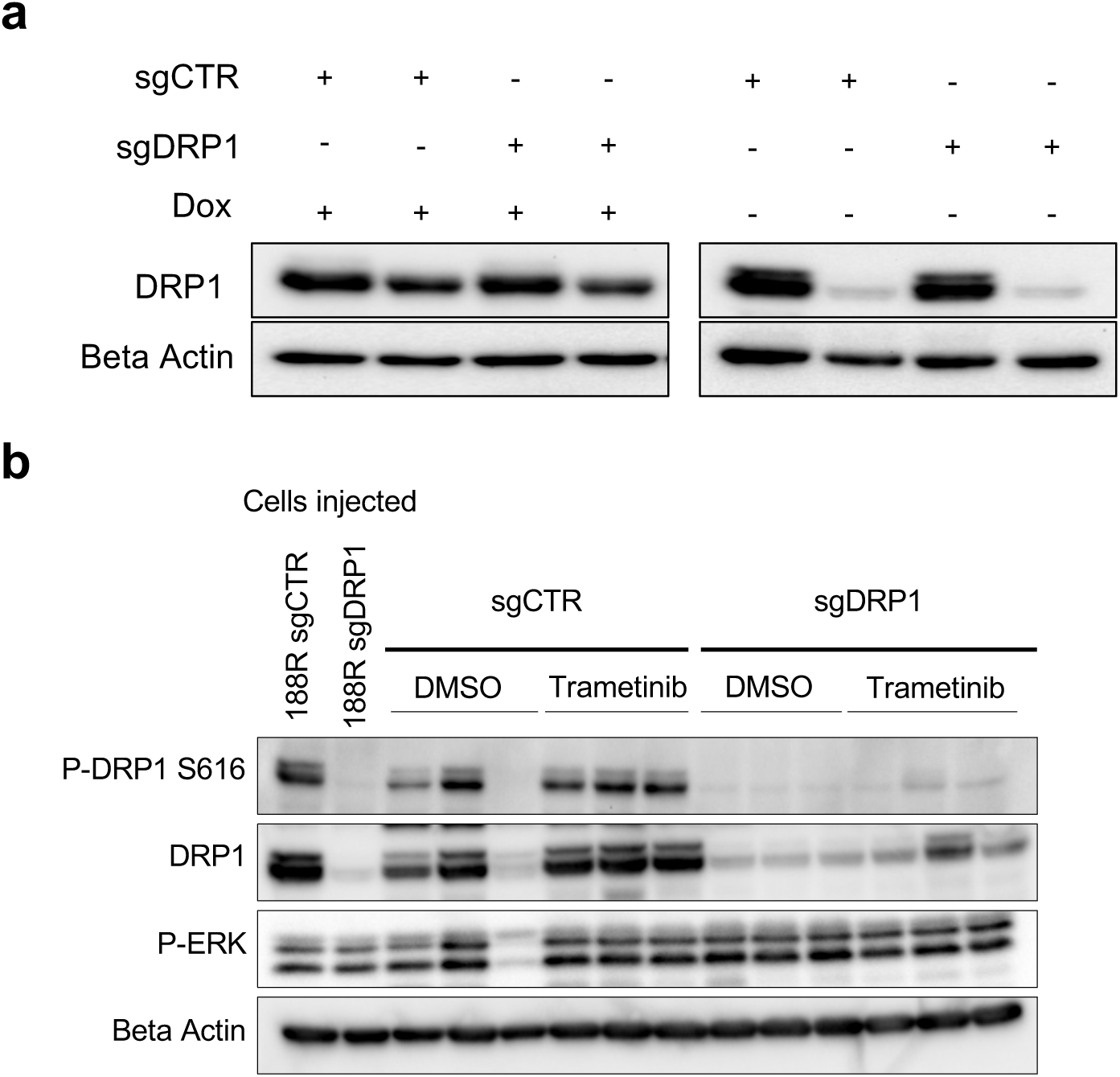
(a) 188R cells were engineered to express doxycycline-inducible mouse DRP1 followed by either sgCTR or sgDRP1 to knock out endogenous DRP1. Removal of doxycycline leads to loss of mouse DRP1 expression and acute DRP1 knockout. (b) Western blot analysis of lysates generated from the tumors harvested from the experiment shown in Fig. 6g.

## Notes

### Competing Interest Statement

The authors have declared no competing interest.

